# Activated MST2 kinase is free of kinetic regulation

**DOI:** 10.1101/2022.01.13.476221

**Authors:** Thomas J. Koehler, Thao Tran, Jennifer M. Kavran

**Affiliations:** Department of Biochemistry and Molecular Biology, Bloomberg School of Public Health, Johns Hopkins University, Baltimore, Maryland, 21205; Department of Biophysics and Biophysical Chemistry, School of Medicine, Johns Hopkins University, Baltimore, Maryland, 21205; Department of Oncology, School of Medicine, Johns Hopkins University, Baltimore, Maryland, 21205

**Author notes:** These authors contributed equally (Thomas J. Koehler, Thao Tran). **Corresponding Author** To whom correspondence should be addressed: Jennifer M. Kavran, Department of Biochemistry and Molecular Biology, Bloomberg School of Public Health, Johns Hopkins University, 615 N. Wolfe Street, Baltimore, MD 21205,.

## Abstract

Canonically, MST1/2 functions as a core kinase of the Hippo pathway and non-canonically is both activated during apoptotic signaling and acts in concert with RASSFs in T-cells. Faithful signal transduction relies on both appropriate activation and regulated substrate phosphorylation by the activated kinase. Considerable progress has been made understanding the molecular mechanisms regulating activation of MST1/2 and identifying downstream signaling events. Here we present a kinetic analysis analyzing how the ability of MST1/2 to phosphorylate substrates is regulated. Using a steady state kinetic system, we parse the contribution of different factors including the domains of MST2, phosphorylation, caspase cleavage, and complex formation to MST2 activity. In the unphosphorylated state, we find the SARAH domain stabilizes substrate binding. Phosphorylation, we also determine, drives activation of MST2 and that once activated the kinase domain is free of regulation. The binding partners SAV1, MOB1A, and RASSF5 do not alter the kinetics of phosphorylated MST2. We also show that the caspase cleaved MST2 fragment is as active as full-length suggesting that the linker region of MST2 does not inhibit the catalytic activity of the kinase domain but instead regulates MST2 activity through non-catalytic mechanisms. This kinetic analysis helps establish a framework for interpreting how signaling events, mutations, and post-translational modifications contribute to signaling of MST2 *in vivo*.

## INTRODUCTION

MST1/2 is a member of the Sterile 20-like kinase family involved in multiple signal transduction pathways and comprised of a kinase domain followed by a flexible linker and a SARAH-domain^1^. Most notably, MST1/2 is the first kinase in the core kinase cassette of the Hippo pathway, a tumor-suppressive pathway and, during development, an anti-proliferative pathway. MST1/2 is also activated in response to apoptotic stimuli in FAS mediated signaling and immune regulation^2^. Activation of MST1/2 relies on phosphorylation of the T-loop on residue T180/183 by *trans*-autophosphorylation^3–5^ or, less frequently, phosphorylation by Tao-1^6,7^. For Hippo signaling, events that increase proximity of MST1/2 to itself trigger autophosphorylation and, thus, activation^8^.

Following activation MST1/2 is primed to phosphorylate downstream substrates, and the identity of these subsequent steps have been established. In the Hippo pathway, active MST1/2 phosphorylates other members of the core kinase cassette, including MOB1A and LATS1/2, the kinase directly downstream of MST1/2 in the Hippo cascade^9^. During apoptotic signaling, activated MST1/2 is cleaved by caspase 3 and then translocates to the nucleus and phosphorylates apoptotic specific substrates such as FoxO and H2B^10^. The factors regulating the activity of MST1/2 during these signaling events remains poorly understood.

The other domains, outside of the kinase domain, are likely modulators of MST1/2 activity. The linker region is, in fact, considered an inhibitory domain. Cleavage of MST1/2 by caspase 3 removes both the linker and SARAH domain. MST1/2 fragments, isolated from cells, lacking the linker region displayed both higher levels of activity and different substrate selectivity than the full-length enzyme^10–16^. The SARAH domain promotes autophosphorylation^5,8,11,17–19^ but has not yet been implicated in directly regulating substrate phosphorylation.

Complex formation with other components of the Hippo pathway also regulates MST1/2 activity. The linker region of MST1/2 contains multiple sites of autophosphorylation that form the binding sites for the Hippo pathway component MOB1A^20,21^. MOB1A is integral to Hippo signal transduction, particularly in promoting phosphorylation of the transcription factor YAP^22^. While numerous roles have been postulated for how MOB1A contributes to pathway activity, the two main theories focus on each of the two core kinases either MOB1A binds the LATS1/2 kinase inducing a conformational change that relieves autoinhibition of LATS1/2^20,23–26^ or MOB1A promotes LATS1/S phosphorylation by MST1/2, perhaps by acting as a scaffold between MST1/2 and LATS1/2^20,21,27–29^. Whether or not phosphorylation of the linker domain or complex formation with MOB1A influences either MST1/2 structure or function is unknown.

The SARAH domain of MST1/2 either forms homodimers, which stimulate autophosphorylation, or heterodimers with the SARAH-domains of either SAV1 or RASSF family members, and each of these interactions tune pathway activity^5,17,30,31^. SAV1 promotes Hippo pathway activity in part by contributing to MST1/2 activation by multiple routes -- increasing the effective concentration of MST1/2, translocating it to the membrane, and inhibiting the activity of the STRIPAK phosphatase^8,18,32–35^. Additionally, SAV1 scaffolds MST1/2 to its substrate LATS1/2^36^ but whether this complex contributes to downstream signaling remains to be determined. The effect of RASSF family members on MST1/2 activity is complicated, RASSFs have been shown to both activate and inhibit MST1/2. *In vitro* RASSF5 (also known as Nore1 and RAPL) and RASSF1A inhibit autophosphorylation of MST1/2 by blocking homodimerization of MST1/2 SARAH domains but in cells can stimulate the activity of MST1/2^5,37,38^. This disparity suggests additional events are responsible for modulating MST1/2. Explanations for spectrum of outcomes associated with RASSFs include mediating cross-talk with other signaling pathways^39^, preventing dephosphorylation and, thus, inactivation of MST1/2^40,41^ similar to the role of its *Drosophila* homolog^42^, functional differences between the six RASSF homologs, and inhibition of MST1/2 activity^38^. It was recently demonstrated that the interaction between MST1/2 and RASSFs are not solely mediated by SARAH domains but includes additional domains from each protein^43^ revealing that our current understanding of the nature of this complex as well as the functional outcomes of it are still evolving.

The current picture of the kinetic regulation of MST1/2 is incomplete. Whether or how intra-domain interactions, phosphorylation, and complex formation contributes to MST1/2 regulation remains to be determined. In this work, we used purified proteins to establish a reconstituted kinase assay to directly monitor kinase activity of MST2. Using this steady-state system we performed quantitative enzymology to determine the factors regulating kinetic activity of MST2 and directly parse the contributions of each domain, phosphorylation, and complex formation to kinetic regulation of MST2. We find that following activation loop phosphorylation MST2 is free of regulation by either intra-domain interactions or complex formation with other components of the Hippo pathway (MOB1A, RASSF5, or SAV1). In the unphosphorylated state, we identify a new role for the SARAH-domain in stabilizing interactions with substrates. We also find that caspase cleavage does not have a kinetic effect on MST2 suggesting the linker region is not inhibitory at the kinetic level.

## MATERIALS AND METHODS

### Expression and purification of MST2

Nucleotides corresponding to human MST2-FL (residues 1-484)(UniProtKB Q13188-1), MST2-KL (residues 1-433), MST2-CC (residues 1-322), or MST2-K (residues 1-314) were cloned into a modified pBAD4^44^ downstream from N-terminal hexahistidine and SUMO tags. MST2 variants were co-expressed with MBP-tagged λ-Phosphatase, encoded in pRSF-Duet (EMD-Millipore, MA) in T7 Express cells (New England BioLabs, MA). Cells were grown at 37°C to OD_600_ of 0.8 at which time 0.5 mM Isopropyl β-d-1-thiogalactopyranoside (IPTG) was added to induce protein expression, and then cells were further grown at 20°C overnight. Cells were lysed in 50mM Tris pH8.0, 400mM NaCl supplemented with protease inhibitors (Sigma-Aldrich, MO). Clarified lysates were incubated with nickel-charged Profinity IMAC resin (Biorad, CA) for one hour at 4°C, and protein eluted with 125mM imidazole. Purified protein was incubated with SENP Protease (made in house) to remove affinity tags. Cleaved protein was further purified by anion exchange and size-exclusion chromatographies. The final protein sample was concentrated to approximately 10mg/mL in 20mM Hepes pH 7.5, 200mM NaCl, and 1mM dithiothreitol (DTT) and flash frozen in liquid nitrogen. To generate phosphorylated MST2 variants, following ion exchange but prior to size-exclusion, protein was incubated with 5mM ATP and 10mM MgCl_2_ for 30 minutes at room temperature, and the reaction quenched by addition of 20 mM Ethylenediaminetetraacetic acid (EDTA).

### Expression and purification of MST2 binding partners

Genes encoding the full-length human RASSF5 isoform D (hRASSF5, UniProtKB Q8WWW0-2), the full-length human MOB1A (UniProtKB Q9H8S9-1) with two site specific substitutions T12A and T35A (hMOB1A^T2A^), or the SARAH domain (residues 531-608) of Salvador from Drosophila melanogaster (dSAV-SARAH)(UniProtKB Q9VCR6)were cloned into a modified pBAT vector downstream from H6 and SUMO tags^44^. Each of the three binding proteins were purified using the same basic protocol, as previously described^30^. Each protein was expressed in T7 Express cells (New England BioLabs, MA) grown in Terrific Broth at 37°C following induction with 0.5mM IPTG. Cells were lysed in 50mM Tris pH 8, 400mM NaCl, 10% glycerol. Proteins were purified by Nickel charged Immobilized Metal Affinity Chromatography (BioRad, CA). The affinity tags were removed during overnight incubation with SENP (made in-house), and the untagged protein further purified by gel-filtration chromatography, concentrated to a minimum of 100μM, and flash frozen in liquid nitrogen. Modifications to this protocol included purification and concentration of each protein prior to gel-filtration. hRASSF5 was concentrated by 45% w/v ammonium sulfate precipitation; hMOB1A^T2A^ was purified on anion-exchange chromatography; dSav-SARAH was purified by anion exchange chromatography and then concentrated by 60% w/v ammonium sulfate precipitation.

### Binding reactions

Phosphorylated MST2-FL was coupled to cyanogen bromide resin (Cytiva, Sweden) at 3mg/mL according to manufacturer’s directions. 40 μL binding reactions contained 20μL column volume (CV) of either blank or MST2-FL conjugated resin and 100μM of either dSAV-SARAH, hMOB1^T2A^, hRASSF5, or Bovine Serum Albumin (BSA) (RPI, IL) in 50mM Hepes pH 7.5, 400mM NaCl, 10% glycerol, 1mM DTT, 10mM MgCl_2_ and was incubated for 1 hour at room temperature. Resin was collected, washed three times with 1 mL of 20mM Tris pH8, 600mM NaCl, 1mM DTT, and resuspended in 35 μL 5X SDS-Loading Buffer. Reactions were analyzed by Coomassie stained SDS-PAGE. Gels were imaged using an Odyssey IR Imaging System (LI-COR, NE). Bands were quantified using ImageJ^45^. Band intensities between different gels were normalized to a common loading control, and then bound protein calculated by subtracting the intensity the bands in the lanes with blank resin from the intensities of the bands in lanes with pMST2-FL resin. Data was plotted using Prism9 (Graphpad Software, CA). P-values were determined using unpaired t-test in R-studio version 1.4.1717^46^.

### In vitro kinase assays

Radiometric kinase assays were carried out as previously described^47^. Briefly, 25μL reactions contained 50mM HEPES pH 7.5, 50mM NaCl, 10mM MgCl_2_, 1mM DTT, 10% glycerol, 0.1mM NaF, 0.1mM Na_3_VO_4_, 0.2mg/ml BSA with either fixed or varying concentrations of either the peptide substrate (Biotin-GGGENWYNTLKRKK-NH2) (Genscript) or ATP spiked with 0.8μCi [γ ^32^P]-ATP (Perkin Elmer). The sequence of the peptide was based on a previously determined consensus sequence for the MST family of kinases to which an additional three glycine residues were included after the N-terminal biotin to act as a linker between the tag and target sequence^48^. All kinase reactions were performed at 30°C and quenched by adding 10μL of 100mM EDTA followed by a 10-minute incubation with 10μL of 20mg/mL avidin (Prospec, Israel) at room temperature. The reaction mixture was then transferred to a 30K Omega Nanosep centrifugal concentrator device (PALL, NY) and washed three times with 100μL of wash buffer (0.5M sodium phosphate, pH 7.5, 0.5M NaCl). The membrane unit was placed in 2mL scintillation fluid (RPI, IL) and counted using a liquid scintillation counter (Beckman, CA). The turnover of limiting substrate was less than 10% for all measurements. Each reaction was performed in duplicate, and measurements were within 20%. Data were fit in Prism (GraphPad Software, CA) to a non-linear curve using the Michaelis-Menten equation to obtain apparent *K*_m_ and *k*_cat_ values.

Concentrations of ATP and peptides used for each experiment are listed in Table 2. The linear range of activity for each MST2 variant with respect to either time or enzyme concentration was established in a series of assays with varying reaction times or enzyme concentrations. Each reaction contained the assay buffer supplemented with 500μM cold ATP and 300μM peptide substrate and enzyme concentrations ranged from 0.5-16nM.

## RESULTS

### Steady State Analysis of MST2

We wanted to understand how the kinase activity of MST2 is regulated. We set out to parse the contributions of phosphorylation, each domain of MST2, and binding partners to the ability of MST2 to phosphorylate substrates. First, we expressed and purified three different length variants of MST2 corresponding to full-length (MST2-FL), the kinase-linker (MST2-KL), and kinase domain (MST2-K). Each variant was mono-dispersed on gel-filtration. Proteins were co-expressed with Lambda phosphatase to yield unphosphorylated, inactive enzymes and to produce active, phosphorylated proteins, ATP and Mg^+2^ were added during purification. Activation loop phosphorylation as monitored by Western blot using a phosphor-specific antibody (pT180) (Figure 1). We then determined steady-state kinetic parameters for MST2 variants using a direct radiometric-assay that monitored phosphoryl-transfer of [γ-^32^P]ATP onto a biotinylated peptide substrate (Table 1) and fit the data to the Michaelis-Menten equation. Assay conditions were determined that maintained a linear relationship between enzyme activity with respect to both enzyme concentration and time (Figures 2,3) indicating that the enzymes used neither degraded nor changed phosphorylation state during the reaction.

**Table 1.** Enzymatic parameters of different MST2 variants and MST2 containing complexes.

| Enzyme | $K_m^{app} \text{ATP} (\mu\text{M})$ | $K_m^{app} \text{peptide} (\mu\text{M})$ | $k_{cat} (\text{min}^{-1})$ | $k_{cat}/K_m^{app} \text{ATP} (\mu\text{M}^{-1} \text{min}^{-1})$ |
| --- | --- | --- | --- | --- |
| Unphosphorylated MST2-K | $84 \pm 33^a$ | $625 \pm 335^{a,b}$ | $74 \pm 22^{a,c}$ | $0.9^{a,c}$ |
| Unphosphorylated MST2-KL | $84 \pm 27^a$ | $805 \pm 140^{a,b}$ | $62 \pm 6^{a,c}$ | $0.7^{a,c}$ |
| Unphosphorylated MST2-FL | $195 \pm 44$ | $139 \pm 18$ | $130 \pm 6$ | $0.7$ |
| pMST2-K | $100 \pm 22$ | $27 \pm 4$ | $1640 \pm 54$ | $16.4$ |
| pMST2-KL | $132 \pm 31$ | $39 \pm 6$ | $1958 \pm 72$ | $14.8$ |
| pMST2-FL | $158 \pm 42$ | $32 \pm 3$ | $1570 \pm 34$ | $10.0$ |
| pMST2-CC | $167 \pm 34$ | $41 \pm 4$ | $1801 \pm 42$ | $10.8$ |
| pMST2-FL + BSA | $78 \pm 7$ | $43 \pm 10$ | $1668 \pm 47$ | $21.5$ |
| pMST2-FL + hMOB1A<sup>2TA</sup> | $79 \pm 14$ | $37 \pm 5$ | $1370 \pm 77$ | $17.4$ |
| pMST2-FL + dSav-SARAH | $91 \pm 7$ | $60 \pm 8$ | $2047 \pm 50$ | $22.5$ |
| pMST2-FL + RASSF5 | $76 \pm 10$ | $52 \pm 10$ | $1567 \pm 60$ | $20.6$ |

<sup>a</sup> Saturating conditions of peptide were not possible so values reported are apparent
<sup>b</sup> Values represent lower limits of measurement
<sup>c</sup> Values represent upper limits of measurement.

**Table 2.**
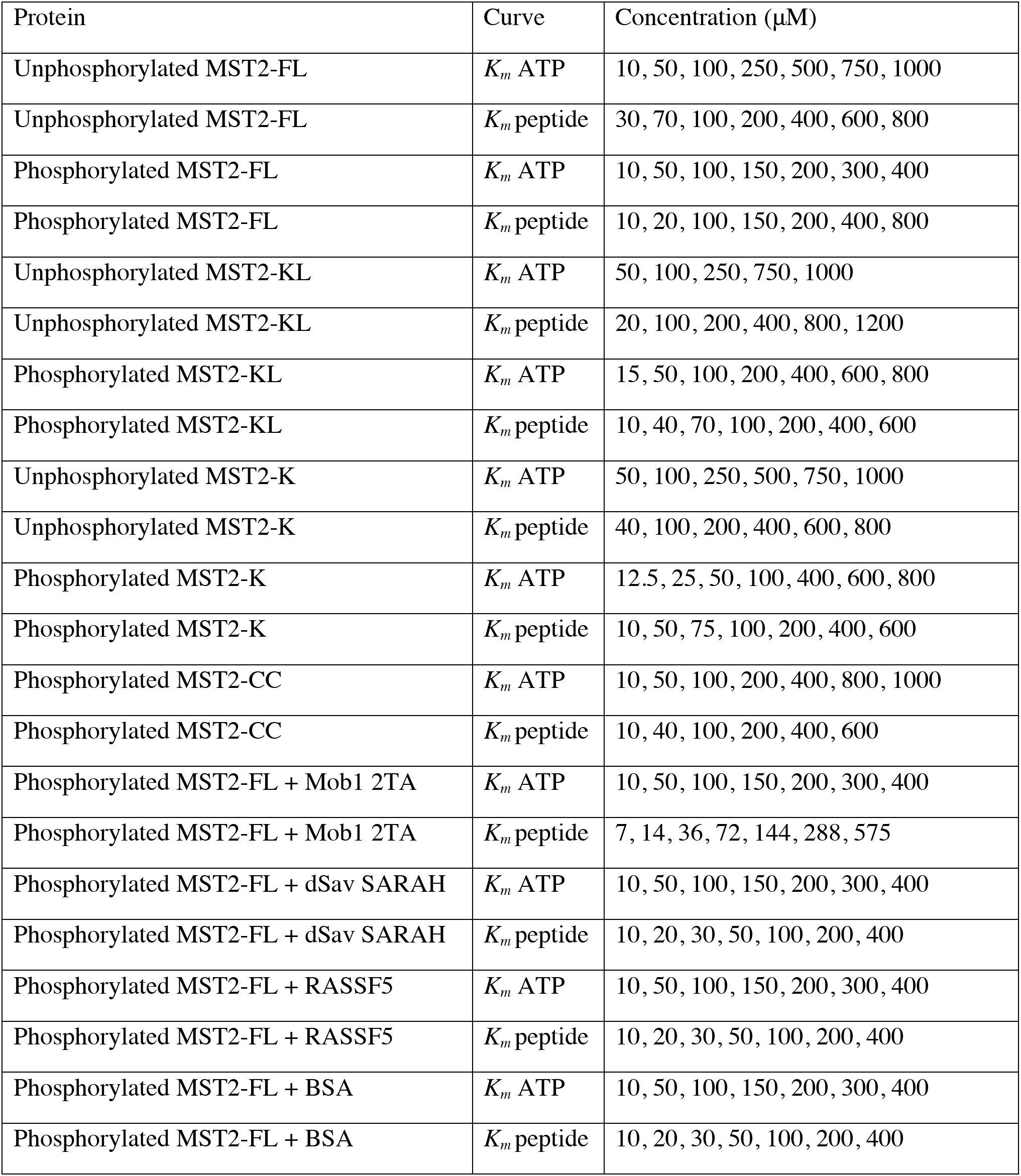
Concentration of ATP or peptide used in Michaelis-Menten experiments.

**Figure 1.**
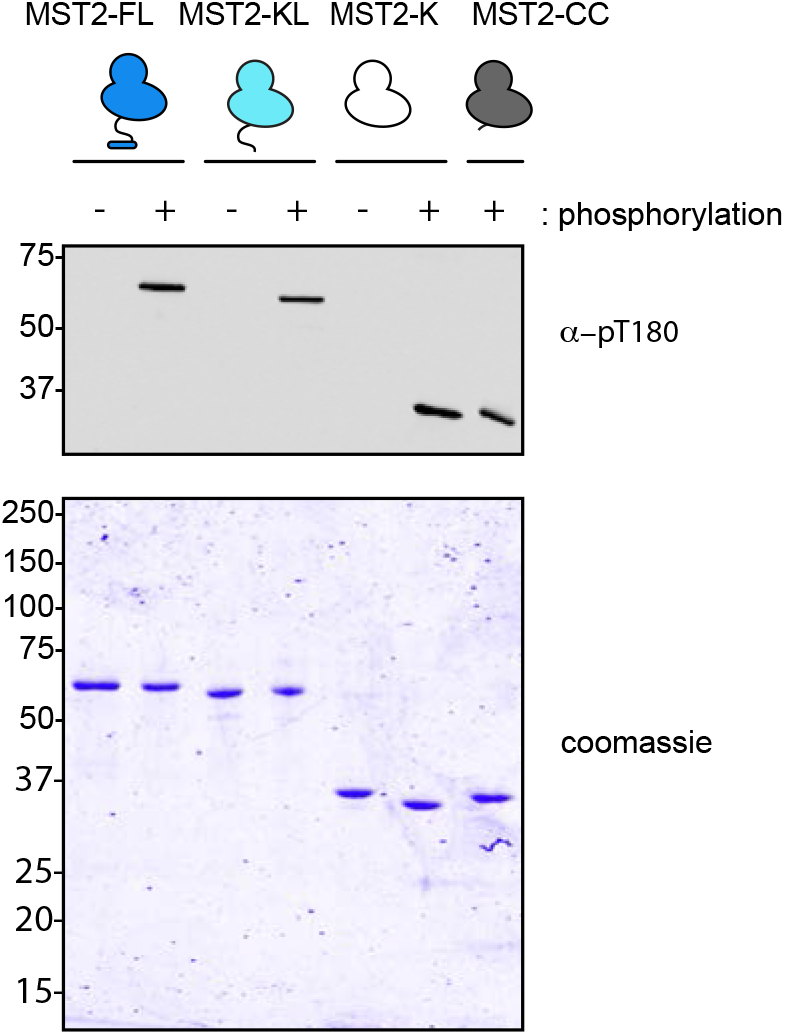
Purification of MST2 variants in defined phosphorylation states. (top) Western blot using a phospho-specific antibody that recognized pT180 on the activation loop or Coomassie stained SDS-PAGE gel (bottom) of MST2 variants including MST2-FL, MST2-KL, MST2-K, and MST2-CC purified in either the unphosphorylated (-) or phosphorylated (+) states.

**Figure 2.**
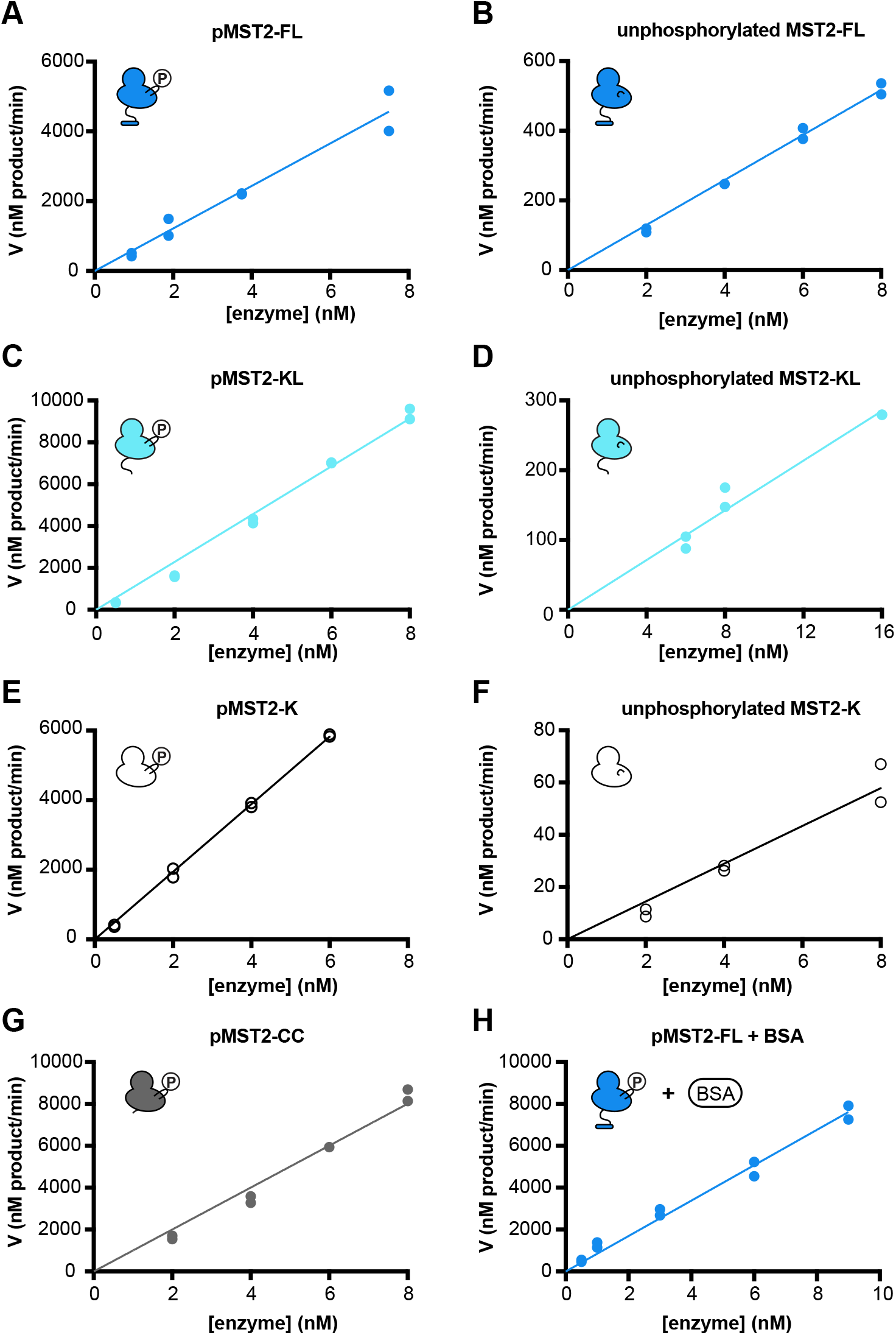
Enzyme behavior of MST2 variants is linear with respect to enzyme concentration. Representative plots showing the linear relationship between velocity (nM of product/min) and enzyme concentration (nM) for both phosphorylated or unphosphorylated MST2-FL (A, B), MST2-KL (C, D), MST-K (E, F), pMST2-CC (G), or pMST2-FL in the presence of BSA (H). Each data point was performed in duplicate and is shown separately.

**Figure 3.**
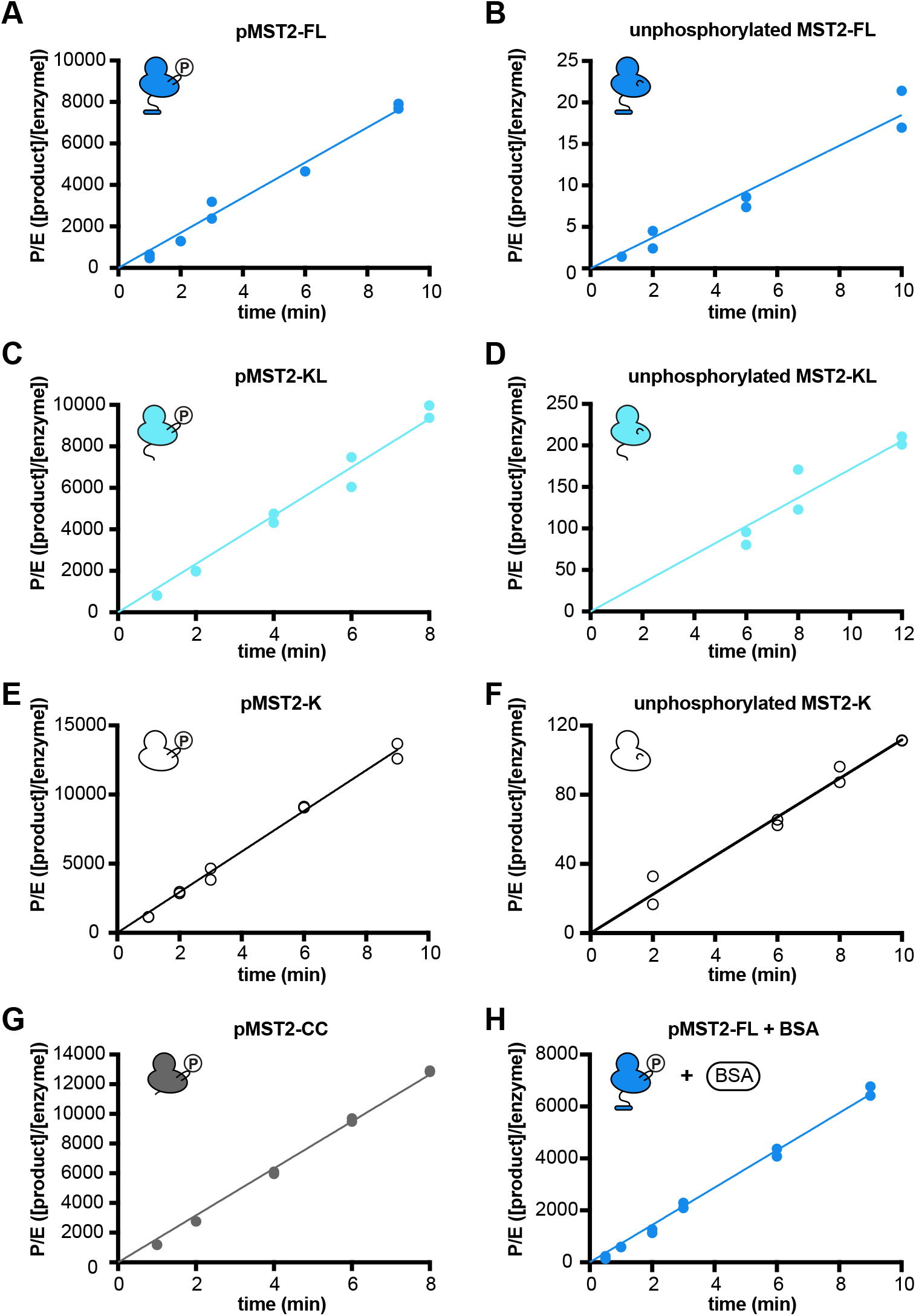
Enzyme behavior of MST2 variants is linear with respect to time. Representative plots showing the linear relationship between [Product]/[Enzyme] with respect to time (minutes) for phosphorylated or unphosphorylated MST2-FL (A, B), MST2-KL (C, D), MST-K (E, F), pMST2-CC (G), or pMST2-FL and BSA (H). Each data point was performed in duplicate and is shown separately.

### The SARAH domain stimulates the activity of unphosphorylated MST2

Unphosphorylated MST2-FL is more active than unphosphorylated variants lacking the SARAH-domain (MST2-KL and MST2-K) (Figure 4, Table 1). These differences are manifested in each kinetic parameter determined but most pronounced in interactions with the peptide substrate. Determination of true *K_m_* and *k*_cat_ values requires addition of saturating amounts of both substrate and ATP to the reaction; owing to the extremely high *K_m_* values for some of the variants tested these conditions were not possible, and so our reported *K_m_* and *k*_cat_ values are considered apparent. The *K_m_*^app^_peptide_ for the variants lacking the SARAH domain, MST2-K (625 μM) and MST2-KL (804 μM), are 3-4 times higher than that of MST2-FL (139 μM). MST2-FL (130min^−1^) is at least twice as fast as variants lacking the SARAH-domain, MST2-KL (62min^−1^) and MST2-K (74min^−1^). Together, these results suggest that the SARAH domain stimulates activity of unphosphorylated MST2. Additionally, we observed a modest decrease (25%) in the *K_m_*^app^_peptide_ between unphosphorylated MST2-KL and unphosphorylated MST2-K suggesting the linker also regulates substrate binding (Figure 4A, Table 1).

**Figure 4.**
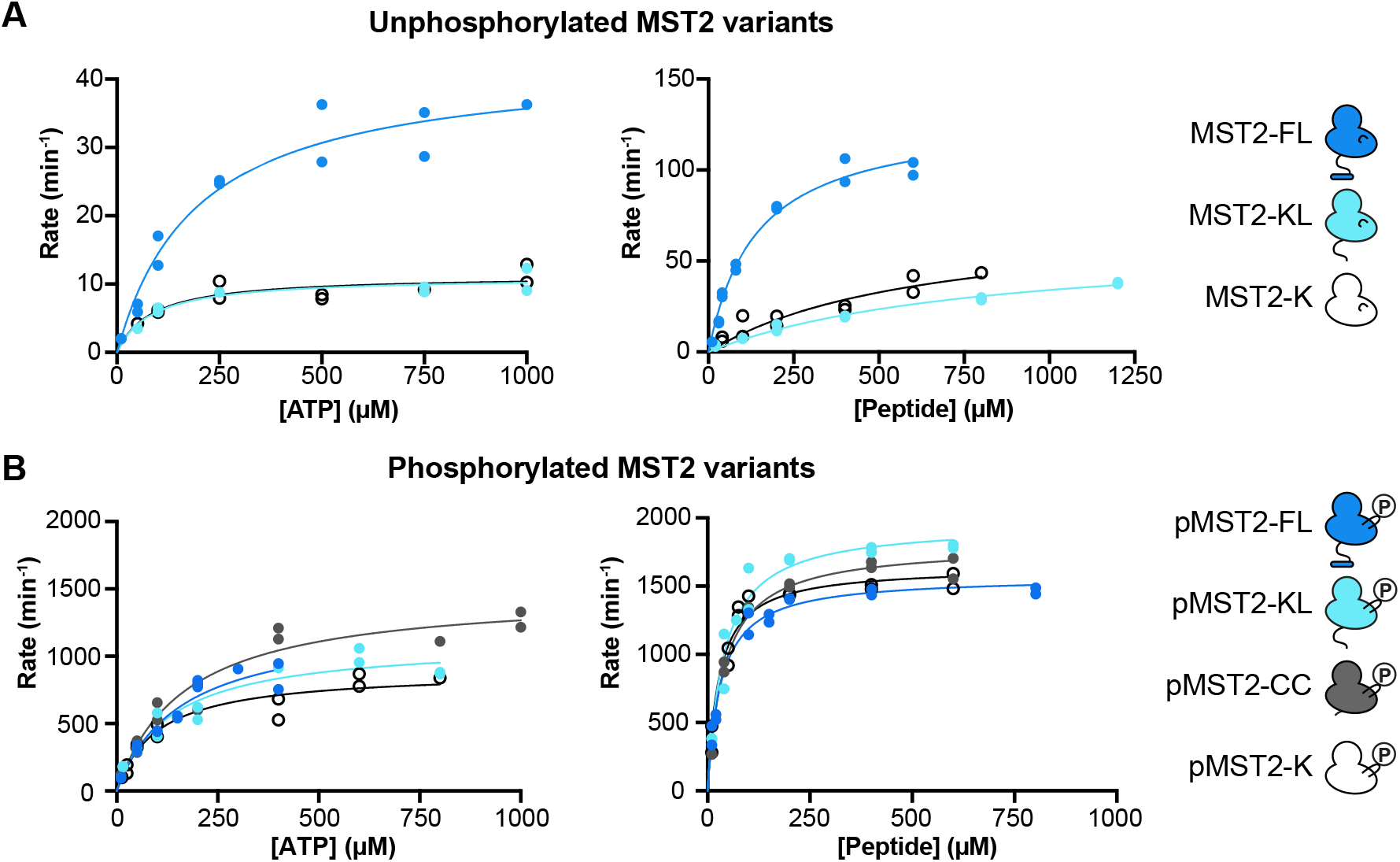
Representative curves of steady-state kinetic analyses for each MST2 variant. Each data point was performed in duplicate and is shown separately for either unphosphorylated (A) or phosphorylated (B) MST2 variants.

### Phosphorylated MST2 is free of internal regulation

Phosphorylated MST2 variants (pMST2-FL, pMST2-KL, and pMST2-K) are similarly active and all are markedly more active than their unphosphorylated counterparts (Figure 4, Table 1). The largely similar activities for the entire set of phosphorylated variants suggests following phosphorylation of the activation loop the activity of MST2 is no longer tuned by either the linker or SARAH-domain. For all the variants investigated, phosphorylation increased turnover, decreased *K_m_*^app^_peptide_, and did not affect *K_m_*^app^_ATP_. The magnitude of these changes, however, differed between variants. The activity of MST2 variants lacking the SARAH-domain were more dramatically stimulated following phosphorylation than MST2-FL suggesting that however the SARAH-domain stabilized interactions of the unphosphorylated kinase domain with peptide substrate, that stabilization is superseded by activation loop phosphorylation. Due to the limitations associated with measuring the *K_m_*^app^_ATP_ for unphosphorylated MST2-KL and MST-K, we cannot make any conclusions on how phosphorylation affected their interaction with ATP.

### Caspase cleavage of MST2 does not alter MST2 activity

Caspase cleavage of MST2 truncates MST2 8 amino acids beyond the kinase domain^13^. In cells, truncated MST2 has elevated kinase activity compared to MST2-FL, perhaps a consequence of removing the autoinhibitory linker domain. Our investigations, however, revealed minimal differences between the activity of pMST2-FL and pMST2-K. Perplexed by our results, we directly monitored activity of a MST2 variant that exactly corresponds to the phosphorylated caspase-cleavage product (pMST2-CC) (Table 1, Figure 4). The catalytic efficiency of pMST2-CC is within 10% of pMST2-FL suggesting that the contribution of caspase cleavage to MST2 activity does not result from intrinsic alterations to MST2 kinase catalytic activity.

### Activity of MST2 is unaffected by binding partners

We also wanted to understand whether or how complex formation with components of the Hippo pathway affected the kinase activity of MST2 by repeating our radiometric assays using pMST2-FL in the presence of hRASSF5, hMOB1A, and SAV1^5,32,49^. First, we expressed and purified each binding partner starting with full-length hRASSF5. Since MOB1A is a substrate of MST2 and its phosphorylation weakens interactions with MST2^20,21,27^, to ensure MOB1A remained unphosphorylated during the course of the reaction we used a variant in which both phosphorylation sites were mutated to alanine (hMOB1A^T2A^). Attempts to purify either full-length SAV1 or the SARAH-domain of SAV1 were unsuccessful, instead we used the SARAH domain from the *Drosophila* homolog (dSAV-SARAH). *Drosophila* Salvador shares both sequence, structural, and functional similarity with SAV1, and previous studies of mixed systems (*Drosophila* Salvador and human MST2) have displayed similar behavior as equivalent mammalian systems^8,18,30,32,35,50^. Then, we confirmed that each purified variant did indeed form a complex with pMST2-FL using an *in vitro* pull-down assay (Figure 5).

**Figure 5.**
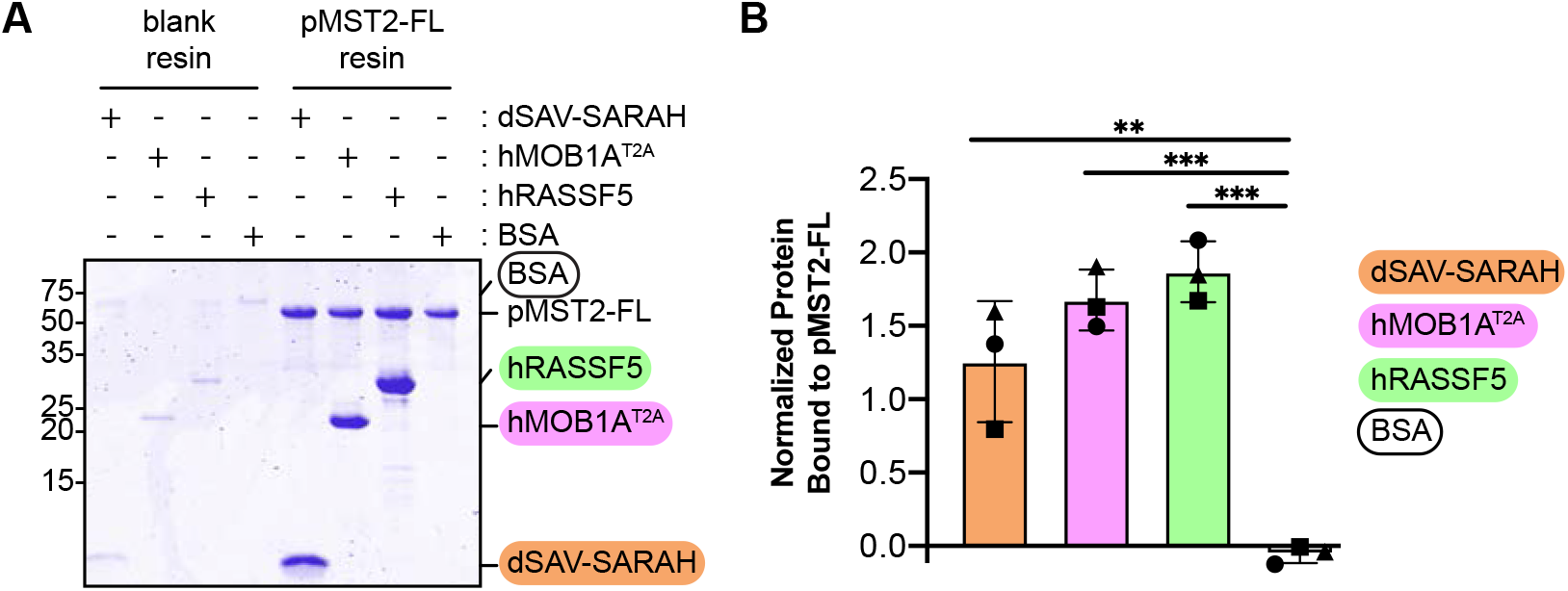
Complex formation of MST2-FL with binding partners. Mixtures containing either dSAV-SARAH (orange), hMOB1A^T2A^ (pink), hRASSF5 (green) or BSA (white) were incubated with either blank or pMST2-FL coupled cyanogen bromide resin. Complexes were isolated, and the amount of protein bound was analyzed by Coomassie-stained SDS-PAGE (A). Reactions were performed in triplicate. The average value of normalized fraction of bound protein is graphed; Individual data points for each replicate are shape-coded; Error bars represent standard deviation. Significant differences were calculated using an unpaired t-test; p-vales corresponding to ≤0.01 (**), or ≤0.001 (***).

We aimed to repeat our kinetic analysis of pMST2-FL in the presence of 100μM of each binding partner, a concentration at least 10-fold over reported disassociation constants to ensure near complete complex formation^20,21,27,43^. Addition of an excess of additional protein could result in macromolecular crowding and alter excluded volumes, diffusion rates, or viscosity; each of these outcomes could manifest as an apparent change in the kinetics of MST2^51^. To first control for macromolecular crowding, we performed our kinetic analysis of pMST2-FL in the presence of an equivalent mass amount of BSA (Table 1, Figure 6). Indeed, we found that while excess BSA did not alter either *K_m_*^app^_peptide_ or *k*_cat_, it did decrease *K_m_^app^*_ATP_ by two-fold. The activity of pMST2-FL in the presence of BSA, therefore, became the baseline we used to compare the effects of binding partners on MST2 activity. We then found that addition of binding partners (hRASSF5, hMOB1A^T2A^, or dSAV-SARAH, hMOB1A^T2A^) did not further tune the activity of pMST2-FL. In each condition tested, the kinetic parameters are within error of the pMST2-FL in the presence of BSA. Our results are in agreement with a previous, qualitative study on the contribution of RASSF5 on pMST2-FL^37^.

**Figure 6.**
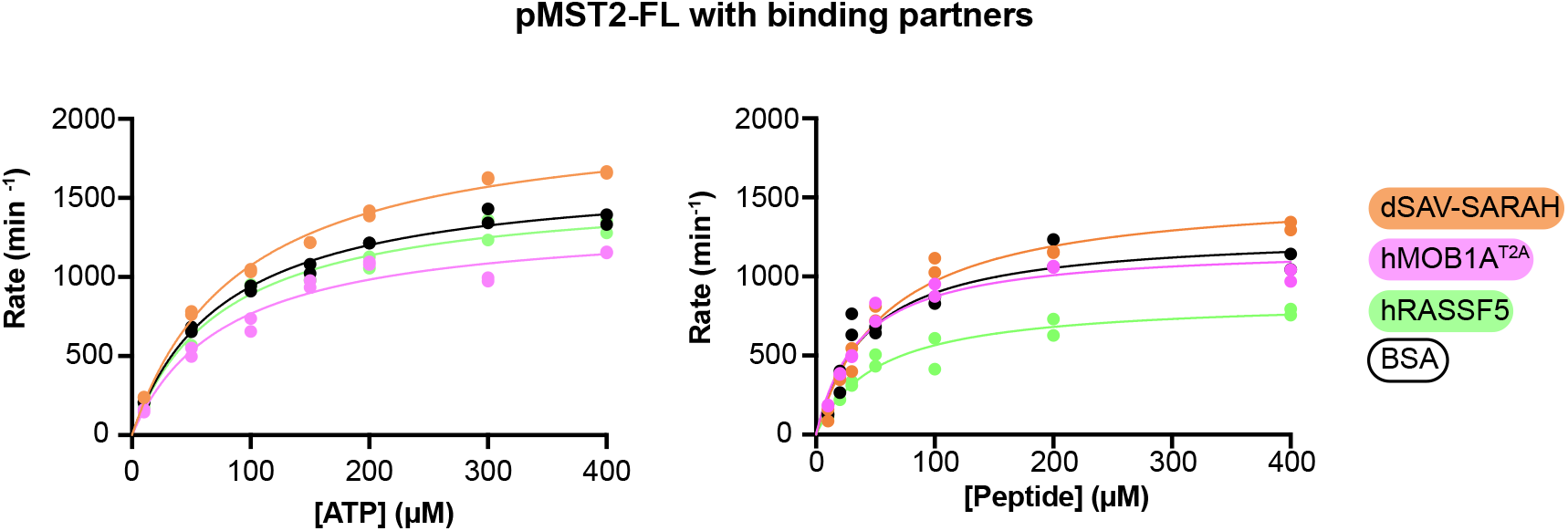
Representative curves of steady-state kinetic analyses for MST2 in the presence of binding partners. Each data point was performed in duplicate and is shown separately for pMST2-FL in the presence of either dSAV-SARAH (orange), hMOB1A^T2A^ (pink), hRASSF5 (green), or BSA (black).

## DISCUSSION

While the mechanism of activation of MST1/2 is relatively well established, how kinase activity is further regulated is less so. Our kinetic analysis reveals that the main determinant of MST2 activity is its phosphorylation state and that when phosphorylated the kinase domain is largely free of kinetic regulation. This activity is not furthered altered by either by interactions with other members of the Hippo kinase cassette or caspase cleavage. We also uncovered a previously underappreciated role for the SARAH-domain in stabilizing interactions with the peptide substrate for unphosphorylated MST2.

We also determined that the activity of pMST2-FL is not tuned by complex formation with other members of the Hippo pathway. These binding partners do, however, contribute to signal transduction by MST2 in cells implying these changes must be attributed to non-kinetic factors. All three binding partners examined mediate larger assemblies which may influence the cellular localization of MST2 or interactions with additional regulatory proteins that influence downstream signaling without altering the kinase activity of MST1/2. For example, SAV1 increases the level of active MST2 by promoting activation and preventing dephosphorylation of active MST1/2, thus promoting downstream signal transduction^8,18,33,34^; hMOB1A may stabilize a ternary complex with LATS1/2 which would increase the effective concentration of a substrate (LATS1/2) for MST1/2^20,21^; and the RASSFs link MST1/2 to other signaling pathways. All of these mechanisms would entail alterations to Hippo pathway activity without a catalytic change in the activity of activated MST2, and any additional analysis of the contribution of these complexes to downstream studies should focus on these non-catalytic contributions.

We were surprised by the lack of difference in activity between pMST2-K and pMST2-FL since the linker region appears to be an inhibitory domain in cells^11,52^. Since caspase cleavage results in an MST1/2 fragment that is 8 amino acids longer than MST2-K, we checked to see if those additional residues had an effect and found that they too did not alter the catalytic properties of phosphorylated MST2. The increased activity of the caspase-cleaved product most likely arises, therefore, from non-catalytic changes. Three previously suggested theories on the increased activity are consistent with both the present and previous data either following caspase cleavage MST1/2 translocates to the nucleus resulting in co-localization of enzyme with its substrate, caspase cleavage changes the substrate preference of the enzyme, or caspase cleavage reduces the likelihood of dephosphorylation of the activation loop^4,10,12,14–16,53^. Future work will be necessary to characterize regulation of activity of pMST2-CC towards full length substrates.

Methodological differences between the current and previous studies most likely account for the reported differences in activity between full-length and caspase cleaved MST2. Of note, our current study uses purified proteins in defined phosphorylation states rather than proteins isolated from cells that most likely have incomplete and varying levels of phosphorylation and uses peptide, rather than protein, substrates which may miss interactions with the substrate outside of the catalytic cleft. Additionally, much of the early work on the inhibitory nature of the linker domain focused on MST1, so sequence variation between the two homologs may accounts for the differences. However, the caspase-cleaved fragments of MST1 and MST2 are 91% identical, and the differences cluster to the αK-helix so it is not readily apparent whether or how these variations would modulate kinase activity.

Comparison of the activities of unphosphorylated and phosphorylated MST2 variants revealed that phosphorylation, expectedly, stimulated activity of MST2. We show that phosphorylation increased turnover, lowered *K_m_*^app^peptide, but did not affect *K_m_*^app^ATP. Phosphorylation improved the catalytic efficiency of MST2-FL by 14-fold and for variants lacking the SARAH-domain at least 18-fold. These trends are in keeping with the common activation mechanism of kinases in which phosphorylation of the activation loop stabilizes the active conformation of the enzyme and improves access of substrates to the catalytic cleft^54^. These trends are consistent with, though greater in magnitude than, previous studies that monitored substrate phosphorylation as an indirect readout of autophosphorylation and activation of MST1^55^.

Analysis of unphosphorylated variants, however, revealed additional roles of the linker and SARAH-domains. At the onset, we wondered if the unphosphorylated linker, which contains multiple potential sites of autophosphorylation, would act as a competitive substrate-inhibitor. The *K_m_*^app^peptide for unphosphorylated MST2-KL is in fact greater than unphosphorylated MST2-K suggesting this is indeed the case, though we note the large error associated with these measurements precludes a definitive analysis. Following phosphorylation, however, the presence or absence of the linker domain does not affect *K_m_*^app^peptide.

Our data also suggests a novel role for the SARAH-domain in stabilizing interactions with peptide substrate. Unphosphorylated MST2-FL has at least a 6-fold lower *K_m_*^app^peptide than variants lacking the SARAH domain (MST2-KL and MST2-K). This additional stabilization is not observed following MST2 activation as all the different length phosphorylated variants had similar values for *K_m_*^app^peptide indicating that the reordering of the active site following activation overrides the contribution of the SARAH-domain in substrate binding. Given the available data, it is unclear how the SARAH-domain contributes to interactions with substrate and structures containing both the kinase and SARAH-domains of MST2 do not provide any additional insight. While minor interactions were observed between the SARAH-domain of RASSF5 with the kinase domain of MST2, similar interactions are not observed in the structure of MST2 bound to SAV1 SARAH-domain^18,37^. Previous discussions of how the SARAH-domain contributes to signal transduction, to this point, have largely focused on how complex formation mediated by the SARAH-domain regulates autophosphorylation and activation of the MST2 but our data suggests that the basal activity of MST2 may be further tuned by the SARAH-domain.

We set out to understand how substrate phosphorylation by MST2 is regulated by performing steady-state kinetics using different length MST2 variants in defined phosphorylation states and in the presence of binding partners. We evaluated the contribution of both internal regulation (phosphorylation or intra-domain interactions) and external factors (protein:protein interactions or caspase cleavage) on substrate phosphorylation by MST2. By using purified recombinant proteins in defined phosphorylation states, we were able to parse the contribution of specific domains, phosphorylation states, and binding partners to regulating the activity of MST2. We find that active loop phosphorylation of MST2 is the largest determinant of its catalytic activity and that following activation MST2 is free of regulation by complex formation with other members of the Hippo pathway, and likely other binding partners as well. Surprisingly we find that MST2-CC is not more active than MST2-FL and, perhaps, that the SARAH-domain can tune the basal activity of unphosphorylated MST2. While there is still much to discern about the contributions of MST1/2 to the complicated signaling landscape of cells, the work presented here will provide key constraints to help interpret how signal transduction is tuned in different cellular contexts.

## DATA AVAILABILITY

All processed data are included in this manuscript. Requests for raw data, further information, or reagents contained within the manuscript are available upon request from the corresponding author.

## ACCESSION CODES

MST2: Q13188-1

RASSF5: Q8WWW0-2

MOB1A: Q9H8S9-1

dSav: Q9VCR6

## AUTOR INFORMATION

### Author Contributions

JMK and TT conceived of this study. JMK, TT, TK conceived and planned experiments. TT and TK performed and analyzed experiments. TT, TK and JMK prepared figures and wrote the manuscript. The manuscript was edited by all authors. All authors have given approval to the final version of the manuscript.

### Funding Sources

This work is supported by NIH R01GM134000 to JMK, NIH T32CA009110 for TT, and NIH T32 GM080189 for TJK.

## Acknowledgements

We thank David Snead and Phil Cole for helpful scientific discussions; Kyler Weingartner and Nam Chu for technical assistance, and Scott Bailey for his use of lab space.

